# Governance networks around grasslands with contrasting management history

**DOI:** 10.1101/2020.03.22.002352

**Authors:** Steluta Manolache, Andreea Nita, Tibor Hartel, Iulia Viorica Miu, Cristiana Maria Ciocanea, Laurentiu Rozylowicz

## Abstract

Romanian grasslands have high nature value, being among the most important biodiversity hotspots at the European level. The European Union Common Agricultural Policy (CAP) contradicts the Biodiversity Strategy to 2020 objective by hindering coordinated grassland governance and collaboration among the involved actors. At the European level, few attempts have been made in creating conceptual strategies for implementing conservation measures in a multi-actor and multi-scale governance setting. Our paper focuses on a comparative network analysis of grassland landscape governance of three Romanian regions (Iron Gates Natural Park – SW; Sighisoara - Tarnava Mare – center; and Dobrogea - SE), representatives for grassland management in mountain and lowland settings. We investigated the structural characteristics of one-mode directed grassland governance networks in the three protected areas (standard cohesion and reciprocity metrics and exponential random graph models), the position of institutions participating in networks (node-level centrality metrics), and the perception of CAP influence on grassland governance by farmers benefiting of CAP agri-environmental payments. In Sighisoara, grasslands governance has been centralized but biodiversity-friendly, while in Iron Gates, grasslands were traditionally managed through a decentralized, community-level system, and this type of governance continues to date. Whereas for Dobrogea’s grasslands, the governance was performed in an intensive, centralized state-run management regime during the communist time and by large landowners after the transition period ended. Our findings illustrate the structure of the three governance networks and dissimilar patterns of collaboration, indicating distinct particularities to be considered when exploring barriers to and options for successful governance in traditionally managed grasslands in the context of CAP measures-driven management.

## 1. Introduction

Semi-natural grasslands of Europe have high economic, socio-cultural, and natural values (Plieninger et al., 2015a). Such landscapes influenced local economies and communities (Bergmeier et al., 2010) and favored the emergence of informal actors that shaped particular norms, rules, knowledge, and skills (O’Farrell and Anderson, 2010). Social demand for agricultural and timber products and economic development of the agricultural and forestry sectors triggered the shift from the traditional multifunctional management providing multiple ecosystem services towards a monofunctional, intensive, management of grasslands focused on food production (Hartel et al., 2015). As a result of this shift, the importance of the traditional management of the grasslands by means of local norms, rules, knowledge, and skills were weakened by the state-enforced regulations such as the EU Common Agricultural Policy (Sutcliffe et al. 2013; Hartel et al., 2020). The prevalence of formal institutional governance coupled with the monofunctional grassland management resulted in a sharp decline of hay meadows and pastures with high socio-cultural and natural values across Europe, especially in the economically developed countries (Overmars et al., 2013). Despite the declining trends in the European continent, traditionally managed grasslands are still present in countries where traditional management practices persist mainly as a legacy of the geopolitical instability (Plieninger et al., 2015b). Europe’s last traditionally managed grasslands provide a unique opportunity to understand the institutional network involved in the governance of these systems mainly because contradictions exist between the 2014-2020 Common Agricultural Policy (CAP) and the European Union (EU) Biodiversity Strategy to 2020 (Balázsi, 2018; Berg et al. 2019). These contradictions also create conflicts between the users of these grasslands (Milczarek-Andrzejewska et al., 2018). This situation can be interpreted as a case of “rigidity trap” (Carpenter and Brock, 2008), where the landowners recognize the unsuitability of the CAP measures for the grassland social-ecological system (Leventon et al., 2017) but are nevertheless encouraged to implement even undesired practices in order to receive EU payments. Furthermore, the acquisition of agricultural lands by large landowners (land concentration) allows the control of large swaths of land by a small number of owners (Calota and Patru-Stupariu, 2019; Constantin et al., 2017). Such threats may simplify the management of grasslands by promoting monofunctionality, change traditional connections, and disrupt traditional land use (Balázsi, 2018; Hartel and Plieninger, 2014).

The governance structures involved in landscape management are often fragmented because policies and operational responsibilities are divided between an array of competing organizations and individuals, often with contradictory objectives (Gieseke, 2019). This is also the case of Romania, for example, in protected areas and water resource governance (Manolache et al., 2018; Nita et al., 2018; Vinke-de Kruijf et al., 2013). The complex governance arenas make the analysis of stakeholders’ interaction and response to policies difficult to understand with conventional tools, and network analysis may provide an ideal framework to acknowledge the formation and coordination of management structures and to provide feedback to policymakers (Bodin et al., 2019; Nita et al., 2018).

Considering the complex socio-cultural and institutional background around multifunctional grasslands, our goal is to employ social network methods to understand governance systems around the management of grasslands from Romania and provide insights for governance strategies of these systems. Network analysis has become an increasingly used approach among practitioners from various research areas: collaborative public management, resource governance, ecology, conservation planning, environmental policy (Berardo et al., 2014; Bodin et al., 2016; Manolache et al., 2018). The investigation of governance networks allows the analysis of informal and formal arrangements where people or organizations work together towards a common goal (Bodin et al., 2019). We refer to individuals, actors and organizations performing repetitive and structured interactions with the aim of managing grasslands at all spatial scales as informal stakeholders (i.e., individuals or groups acting on the base of socially-shared rules, outside of officially-sanctioned channels such as farmers, farmers associations, NGOs) and formal stakeholders or actors (i.e., official entities such as city halls, public agencies, public authorities) (Helmke and Levitsky, 2004; Ostrom, 2005).

We focus on a comparative investigation of network structures around the multifunctional grasslands from three regions of Romania with contrasting management history: Macin Mountains with large communist collective farms, Sighisoara-Tarnava Mare with a mixture of collective and private small farms, and Iron Gates with small private farms. All three regions are covered today by protected areas and are under the CAP influence. In examining the influence of CAP on grassland governance networks we tested the following research hypotheses: 1) The grassland governance networks developed under the CAP influence are shaped by the current administrative context and not past farming practices, expecting more centralized networks in regions with fewer administrative and territorial jurisdictions regardless of past farming management, 2) The EU CAP measures promote local informal stakeholders as network leaders, and 3) The past farming management experience influence the ways how farmers benefiting of agri-environmental payments percept the main issues around grassland management under the CAP influence.

## 2. Methods

### 2.1. Case studies

We selected three contrasting regions as case studies capturing different environments and past farming practices, as follows: a) the region of Macin Mountains (Macin Mountains National Park, hereafter Macin Mountains) within Dobrogea – South-East Romania, representative for grassland management in lowland areas, b) The Saxon cultural region of Central Romania (the area covered by Sighisoara-Tarnava Mare Natura 2000 site, hereafter Sighisoara area, the town of Sighisoara is situated in the central area of the region) which is representative for hilly (tableland) areas and c) The cultural region of Banat (the area covered by Iron Gates Natural Park, hereafter Iron Gates) from South-Western Romania, representative for low elevation mountains areas (Supplementary Data 1).

A centralized, state-run management regime characterized Macin Mountains (Dobrogea) during the communist time. After the transition period ended, Macin Mountains grasslands were managed primarily by large landowners with large collective farms, dairy farms, with little ownership sense (Anthony and Moldovan, 2010). Sighisoara area is known as a tableland with diverse land-use and a mixture of communal and private small farms (Fischer et al., 2012; Sutcliffe et al., 2013). In contrast, Iron Gates from South-West Romania is situated at the border with Serbia, characterized by low altitude mountains, small private farms with diverse land-use (e.g., small patches of corn plantation, hayfields, small orchards). In Iron Gates, grassland management was traditionally performed in a decentralized, community-level system (Pătroescu and Rozylowicz, 2000), and this type of management continues to this day (Manea, 2003). The three protected areas cover one county and 18 local administrative units in Macin Mountains, two counties and 15 local administrative units in Iron Gates, and three counties and 14 local administrative units in the Sighisoara area (Supplementary Data 2).

### 2.2. Network data

We collected network data by face-to-face semi-structured interviews applied in 2018 with representatives of institutions potentially involved in the management of grassland in the three protected areas. To select the interviewed institutions, we reviewed Romanian legal norms and CAP technical documents related to grasslands and extracted local, regional, and national likely to be involved in the governance of grasslands in the three protected areas. This step resulted in a predefined list of stakeholders for each case study. If the interviewed persons nominated actors not included in the predefined list, we applied the interviews to one or two key representatives of these stakeholders. We targeted stakeholders involved in contribution to information flow in grassland management (e.g., informing about direct payments requirements, grassland management technologies, grassland biodiversity, and conservation, funding availability, agri-environment schemes), control of grassland management (e.g., complying with requirements of direct payments, complying with various legal norms), approval of direct payments (submission of direct payments documentation, payments to farmers). The list of interviewed stakeholders included local, regional, and national agencies and authorities, local and regional administrations, private and public enterprises, NGOs, local associative organizations, resource managers, active in grassland management. We did not interview individual farmers and national agencies. The farmers were included in the resulted network as a collective actor. National agencies and authorities were not interviewed because they have only a top-down/indirect influence on the network, their competencies being delegated to local or regional representatives already included in the network (e.g., Ministry of Agriculture to the local Agencies for Payments and Intervention in Agriculture).

The respondents were asked to indicate other stakeholders with whom they were collaborating, the direction of the collaboration (e.g., they send information to the named institution, they receive information from the named institution or both), the type of exchanged information, and the frequency of contact. We considered as exchanged information any regulatory/administrative task, joint projects, participation in the same working groups, drafting reports and studies in common, other shared activities. The semi-structured interviews also included a question about the quality of collaboration on a Likert 5 points scale. However, the representatives of interviewed organizations were reluctant to provide an answer; thus, we discarded these data from our analysis.

With the data gathered for each protected area, we constructed a grassland governance network, where stakeholders (vertices) were linked (through edges) if they transfer any information related to grassland management. We included in the analysis the direction of the information transfer (sender, receiver) and obtained one-mode directed networks (Barabási, 2016; Borgatti et al., 2017). Furthermore, the stakeholders included in the three networks we classified by entity type as follow: (1) public authorities (ministry, local, regional, and national governmental agencies), (2) public administration (city halls), (3) NGOs, and research and education bodies, and (4) enterprises (i.e., industry, private businesses, state-owned businesses, farmers). Public authorities, research and education bodies, industry, private businesses, state-owned businesses act based on a formal mandate, whereas farmers and NGOs based on an informal mandate.

### 2.3. Network-level analyses

To investigate if the structural characteristics of grassland governance networks in the three different protected areas are influenced by farming management history and local administrative complexity, we used standard cohesion and reciprocity metrics, which along with exponential random graph models for one-mode directed network (Barabási, 2016; Borgatti et al., 2017; Lusher et al., 2013) will help to determine the differences between the three contrasting areas analyzed, especially regarding their network structure.

The *cohesion and reciprocity metrics* for directed networks considered here were: size, volume, average degree, network density, network fragmentation, network diameter, average shortest path, network compactness, and arc reciprocity (Borgatti et al., 2017). These metrics allow us to infer which case study displays a higher level of interaction between the actors and an efficient information flow. The *size of the network* is represented by the number of institutions participating in the respective network, while the *network volume* by the number of edges or connections between institutions (Barabási, 2016). In our case, both metrics are influenced by the administrative complexity (e.g., number of counties overlapping the analyzed protected area), protected areas covering more counties being expected to have a larger and more complex governance network. *Network density* represents the proportion of observed connections between institutions participating in transferring information to the maximum number of possible connections in the respective network. This metric indicates the magnitude of interconnectivity between network members (networks with densities closer to 1 are dense, while those with densities closer to 0 are sparse) (Borgatti et al., 2017). *Arc reciprocity* represents the proportion of mutual connections between organizations divided by the total number of connections (reciprocal or otherwise). A network high arc reciprocity indicates a high proportion of bidirectional connections between network actors (Borgatti et al., 2017). The degree of a node (stakeholder in the network) is the number of connections of the respective node regardless of the direction of information transfer. The *average degree* of the network can be used to compare networks, e.g., networks with low density and relatively high average degree indicate a centralization around a small number of highly connected network members (Barabási, 2016).

Average path length and network diameter can be used to assess the information flow in the network. *Average path length* represents the average number of institutions that must be passed to connect two institutions in the network, and the *network diameter* the shortest path length between the two most distant nodes in the network. Information flows faster and more effectively in networks with small average path length, while in networks with small diameter, distant members are well connected (Barabási, 2016). A closely related metric is the *fragmentation* of the network, which measures the proportion of pairs of institutions that cannot reach each other by a path of any length. Higher fragmentation indicates a limited potential for exchanging information (Chen et al., 2007).

We use exponential random graph models (ERGM or p* models) to infer if observed network substructures (configurations) are more commonly observed in the network than expected by chance (Lusher et al., 2013; Robins et al., 2012; Wang et al., 2009). ERGMs are related to general linear models, modified to deal with the lack of independence of observations. More details ERGM statistics and assumptions are provided by Wang et al. (2009). ERGM for directed networks can accommodate several configurations such as density, reciprocity, k-in-stars, k-out-stars, 2-mixed-stars, alternating-in-star (Wang et al., 2009). To fit the ERGM for each governance network, we use P-Net for directed network, as suggested by Lusher et al. (2013): 1) we selected several configurations to be estimated, run the model with default values, and checked the model statistics. Density for all estimations was fixed (i.e., the observed network density); 2) if *t*-ratios for selected network configurations were less than 4 for all values, we updated the parameters with the new estimated values, and re-run the model using an increased multiplication-factor. Because the analysis is trial and error, we repeated the procedure until the model converged or we included/excluded configurations and repeated the first 2 steps if the model was unreliable; 3) if the model converged (all selected parameters had *t*-statistics <0.1 in absolute values) we estimated the Goodness-of-Fit using 100,000,000 simulations. For Goodness-of-Fit analysis, we updated the converged parameters with the values obtained in step 2 and zero for estimates for all other parameters. A model with a good fit has the absolute value of the *t*-ratios <0.1 for converged parameters, and <2 for other network statistics. If the *t*-ratio for any P-Net configuration was over the threshold, we repeated the entire analysis by adding and/or discarding configurations. All configurations from converged models are explained in Table 1, and all configurations in models are visualized in Figure 1.

**Table 1.**
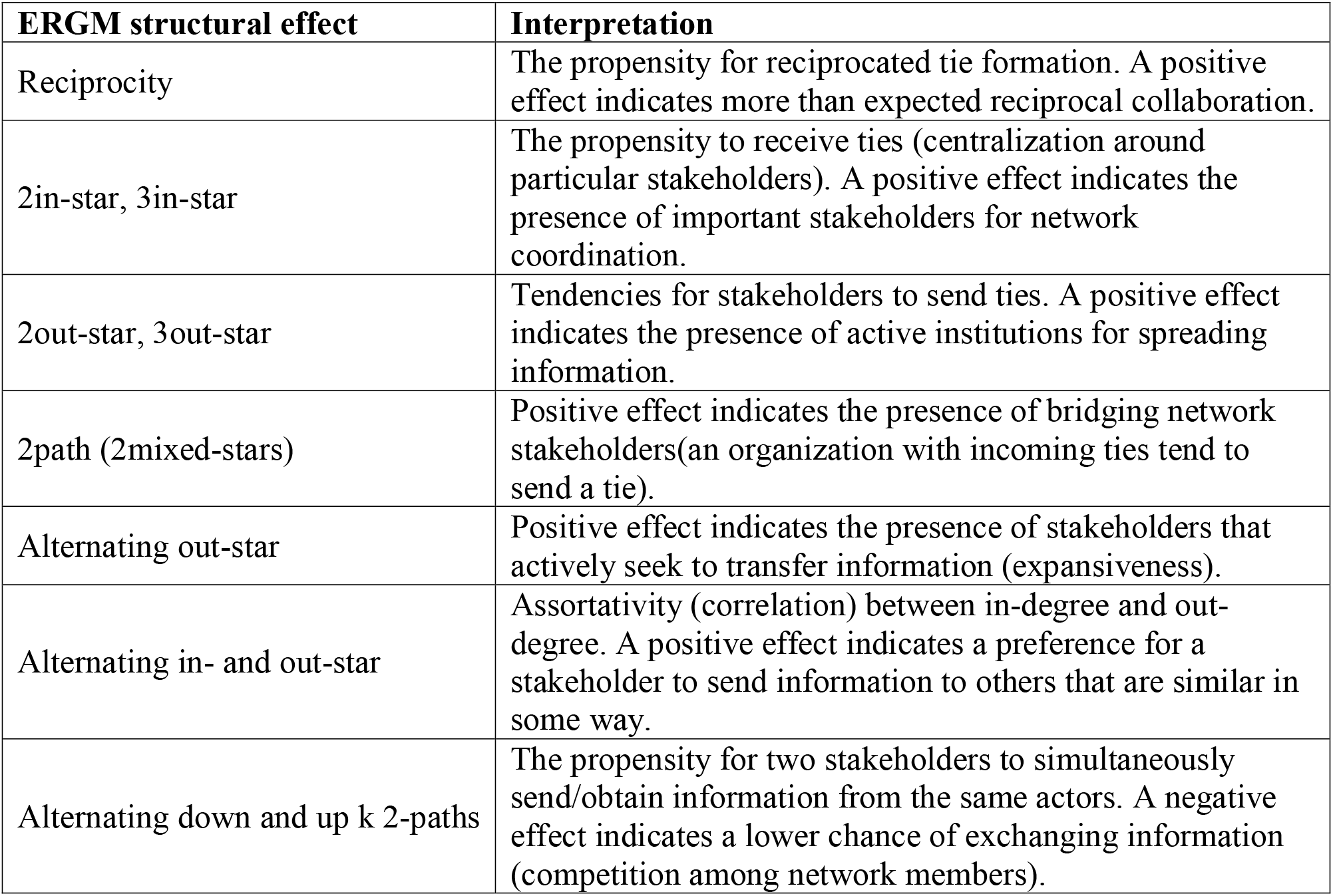
Interpretation of structural effects significant in converged ERGMs (Lusher et al., 2013)

**Figure 1.**
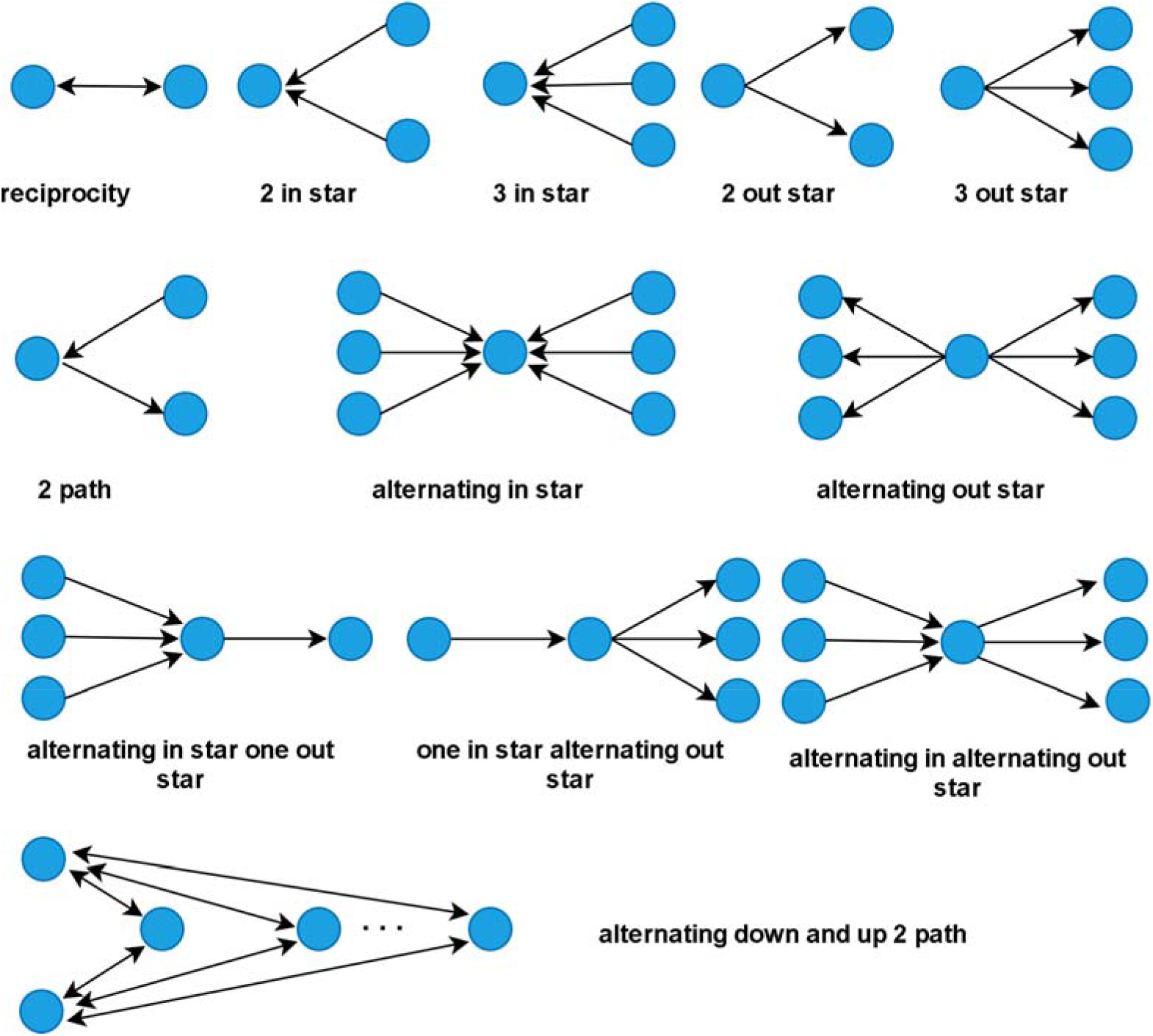
Structural effects (configurations) included in converged ERGMs

### 2.4. Grassland management stakeholders-level centrality metrics

To analyze the position of institutions participating in grassland governance networks within the three protected areas, we calculated the following node-level centrality metrics for one-mode directed networks: normalized in-degree centrality, normalized out-degree centrality, eigenvector centrality, and normalized betweenness centrality (Borgatti et al., 2017). *In*- and *out-degree* centralities represent the number of ties an institution has divided by the maximum number of possible ties in the respective set (incoming, outgoing). In-degree centrality measures the status of the institutions (stakeholders with high in-degree are considered as hubs in resource management, while those with high out-degree are important for initiating management activities (Borgatti et al., 2017). *Betweenness centrality* for an institution is calculated as the number of shortest paths that pass through the respective institution, normalized by the maximum betweenness that any institution in the network can achieve. An actor with a high betweenness score might play an important role in the network because it can mediate the exchange of information between the linked institution (Barabási, 2016). *Eigenvector centrality* calculates the principal eigenvector of the adjacency matrix defining the network, and normalized by dividing the raw eigenvector score by maximum score attainable in the network, capture not only the number of connections but also the influence of neighbor institutions. Because it is a sum of the centrality values of the connected stakeholders, eigenvector measure shows here how central is the actor of interest in the network (e.g., in the best position to transfer information) (Borgatti et al., 2017). Network analyses were performed using Ucinet 6.698 (Borgatti et al., 2002).

### 2.5. Perception of CAP influence on grassland governance

We interviewed farmers from the three protected areas, taking in consideration only those benefiting from one of the following CAP agri-environmental payments (MARD, 2014): a) CAP agri-environmental payments package 1 high nature value grassland, CAP agri-environmental payments package 2 traditional agriculture (only in combination with package 1); b) CAP agri-environmental payments package 2 traditional practices; c) CAP agri-environmental payments package 3 grassland important for birds (*Crex crex*, *Lanius minor*, *Falco vespertinus*). For package 1, the main requirements are indicated by the interdiction to use pesticides and synthetic fertilizers, limited use of manure, delayed haymaking, and low-intensity grazing. For package 2, the applicants must comply with requirements for package 1 and opt for the manual mowing or using small machinery. Package 3 requires a supplement leaving 3 meters of untrimmed grassland (Sutcliffe et al., 2015). The survey administrated to farmers considered their perception of CAP influence on grassland governance and the main issues around grassland management and the links with grassland governance. Therefore, the survey included questions regarding their capacity (owner, administrator) and perception of CAP influence on grassland management (no influence, negative influence, positive influence), and main issues perceived related to grassland management under CAP influence (insufficient funding per ha, late agri-environmental payments, bureaucracy, lack of market for grassland products). The questions were selected after face to face interviews with representatives of Agencies for Payments and Intervention for Agriculture in the three protected areas. We collected 58 responses from farmers as follows: 9 from Macin Mountains, 24 from Sighisoara Tarnava Mare, and 25 from Iron Gates. The answers were analyzed using Multiple Correspondence Analysis (MCA) (Greenacre and Blasius, 2006). MCA allows the evaluation of associations among more than two categorical variables. We performed MCA analyses using FactoMineR and factoextra R packages (Kassambara and Mundt, 2016; Lê et al., 2008). We generated a symmetrically scaled biplot of coordinates of coded answers (active variables) and included protected area and farmers’ capacity as supplementary variables (Greenacre and Blasius, 2006). We considered an answer as a significant contributor to an axis if its eigenvalue exceeds the average of eigenvalues from the respective axis (Kassambara and Mundt, 2016).

## 3. Results

### 3.1. Network-level processes in grassland governance

The size of grassland governance networks varies between 63 (Iron Gates) and 31 organizations (Macin Mountains) (Table 2). In all cases, public authorities and administrative bodies dominate the network. In contrast, public and private enterprises and NGOs and research institutions are only marginally presented (Supplementary Data 3). The grassland governance networks are sparse in terms of densities of connections (Table 2). However, the network within Macin Mountains is the most collaborative (22% of potential connections exist), followed by Iron Gates (12% of potential connections exist) and Sighisoara (8% of connections exist). The three networks are also dissimilar in terms of average connections between participating institutions (average degree). Institutions involved in the grassland management in Iron Gates and Macin Mountains are well linked when compared to Sighisoara (Kruskal-Wallis test in-degree = 23.89, p < 0.001; Kruskal-Wallis test out-degree = 30.21, p <0.001). Macin Mountains and Iron Gates are also similar in terms of reciprocal connections, with over 80% of the ties reciprocated, significantly larger than in Sighisoara, where 65% of connections between involved institutions are in both directions. The average shortest path connecting two institutions (average geodesic distance) is smaller in Macin Mountains than in Iron Gates and Sighisoara. A similar pattern follows the network diameter, i.e., the number of institutions connecting the two most distant connected institutions in the respective network or maximum geodesic distance.

**Table 2.** Cohesion and reciprocity metrics in three grassland governance networks from Romania

| **Network-level metric** | **Sighisoara** | **Iron Gates** | **Macin Mountains** |
| --- | --- | --- | --- |
| Size | 52 | 63 | 31 |
| Average degree | 3.98 | 7.33 | 6.58 |
| Density | 0.08 | 0.12 | 0.22 |
| Fragmentation | 0.15 | 0.12 | 0.12 |
| Diameter | 6 | 5 | 4 |
| Average shortest path | 2.66 | 2.31 | 1.92 |
| Arc reciprocity | 0.65 | 0.81 | 0.84 |

ERGMs with structural effects and fixed propensities for tie formation (network density) converged for all analyzed networks and fitted very well the configurations of observed networks (Table 3, Supplementary Data 4). ERGMs indicate a dissimilar pattern of interactions among the network actors in the three analyzed networks. The propensity for collaborations (reciprocity) is higher than expected by chance in Sighisoara and Macin Mountains but at random in Iron Gates.

**Table 3.** ERGM results for three grassland governance networks from Romania (estimates and standard deviation, and t-ratio). Significant effects (p < 0.05) are in bold and marked with *.

| Effects | Sighisoara |  | Iron Gates |  | Macin Mountains |  |
| --- | --- | --- | --- | --- | --- | --- |
|  | Estimates (sd) | t-ratio | Estimates (sd) | t-ratio | Estimates (sd) | t-ratio |
| reciprocity | **4.35 (0.35)** | **-0.03\*** | -0.23 (0.26) | -0.04 | **4.01 (0.42)** | **-0.03\*** |
| 2-in-star | 0.22 (0.14) | 0.05 | **0.13 (0.05)** | **-0.04\*** | 0.35 (0.29) | -0.04 |
| 2-out-star | **0.68 (0.17)** | **0.03\*** | 0.06 (0.06) | -0.03 | 0.16 (0.33) | -0.06 |
| 3-in-star | -0.003 (0.01) | 0.03 | -0.0003 (0.003) | -0.02 | -0.01 (0.02) | -0.01 |
| 3-out-star | **-0.04 (0.02)** | **0.05\*** | 0.004 (0.01) | -0.01 | -0.003 (0.03) | -0.05 |
| 2path | **-0.16 (0.03)** | **-0.02\*** | 0.01 (0.01) | -0.07 | -0.05 (0.10) | -0.07 |
| AinS | -0.97 (0.58) | 0.01 | -0.39 (0.74) | -0.07 | -1.78 (1.20) | -0.07 |
| AoutS | **-1.54 (0.55)** | **-0.02\*** | -0.36 (0.77) | -0.06 | -1.63 (1.20) | -0.04 |
| Ain1out-star | 0.27 (0.50) | 0.03 | 0.08 (0.08) | 0.05 | 1.62 (1.14) | 0.03 |
| 1inAout-star | 0.82 (0.42) | 0.04 | -0.03 (0.06) | 0.02 | 0.99 (1.21) | -0.01 |
| AinAout-star | **-1.53 (0.60)** | **0.06\*** | -1.09 (0.84) | 0.06 | **-3.00 (1.06)** | **0.07\*** |
| A2P-DU | - | - | **-0.33 (0.04)** | **0.08\*** | - | - |

The dissimilarity of Iron Gates can be explained by the positive 2-in-star effect, which suggests a slightly higher than expected chance for an institution to receive ties from two others (2-in-star centralization) but at random propensity of sending ties to two other institutions (2-out-star popularity). The 2 in star effect, as well as the similar effect one order higher (3-out-star), is positive in Sighisoara, suggesting the presence of popular actors. However, these popular actors do not act as brokers in Sighisoara (receivers of ties also send ties) as 2-path configurations and alternating out stars are lower than expected. Also, alternating out stars is significantly lower than expected in Macin Mountains, suggesting the lack of coordination with more than one institution. ERGM for Iron Gates network converged only by including a higher-order configuration, which a lower chance to exchange information between several actors, confirming the lack of reciprocity at higher orders.

### 3.2. Key stakeholders in grassland governance networks

The network of stakeholders involved in grassland management indicates that the most important actors mediating the transfer of information within the network (betweenness centrality) are mostly public agencies (Figures 2–4, Supplementary Data 3). In Sighisoara area, top institutions are county-level Agencies for Payments and Intervention for Agriculture (in charge with the management of application for agri-environmental payments), farmers and enterprises (as collective actors), and the representative of Ministry for Agriculture in Sibiu county. For Macin, top stakeholders are the county level Agency for Payments and Intervention for Agriculture (in charge with the management of application for agri-environmental payments) and Office for Soil Studies (in charge with soil analyses and classification), followed by protected area administration and the local office of Agency for Payments. In Iron Gates, farmers as collective actors are the main actors mediating the transfer of information, followed by the administration of the protected area and local offices of the Agency for Payments. In Sighisoara and Macin Mountains networks, the number of actors with betweenness centrality of zero (actors with no role in the transfer of information) is higher than in Iron Gates.

**Figure 2.**
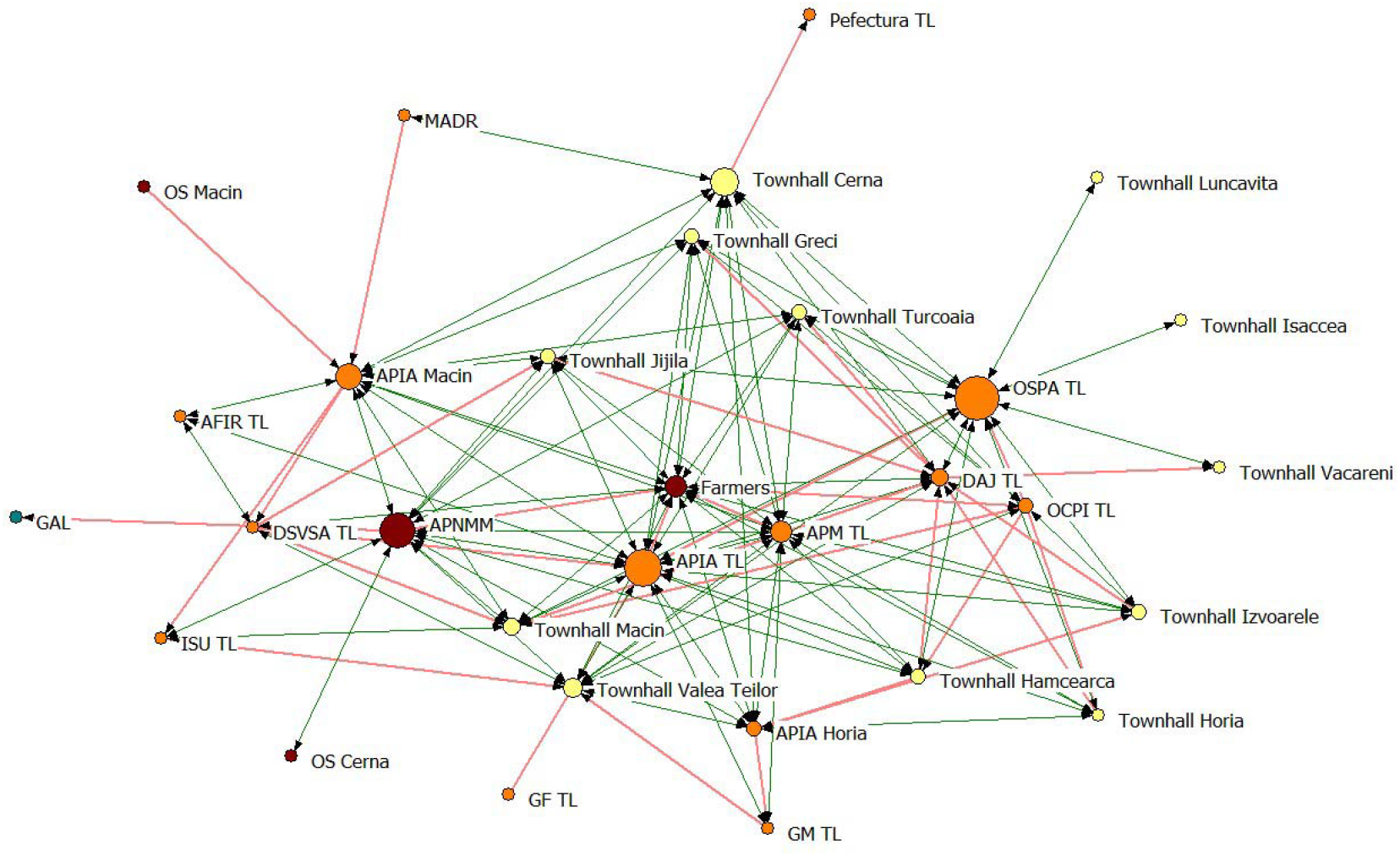
Grassland collaboration network in Macin Mountains (green lines = reciprocal links of collaboration, red lines = one direction of collaboration, orange circles = public authorities, red circle = enterprises, yellow circle = public administration, green circle = NGO & research & education organizations, size of nodes = betweenness centrality). The full names of organizations are available in Supplementary Data 3.

**Figure 3.**
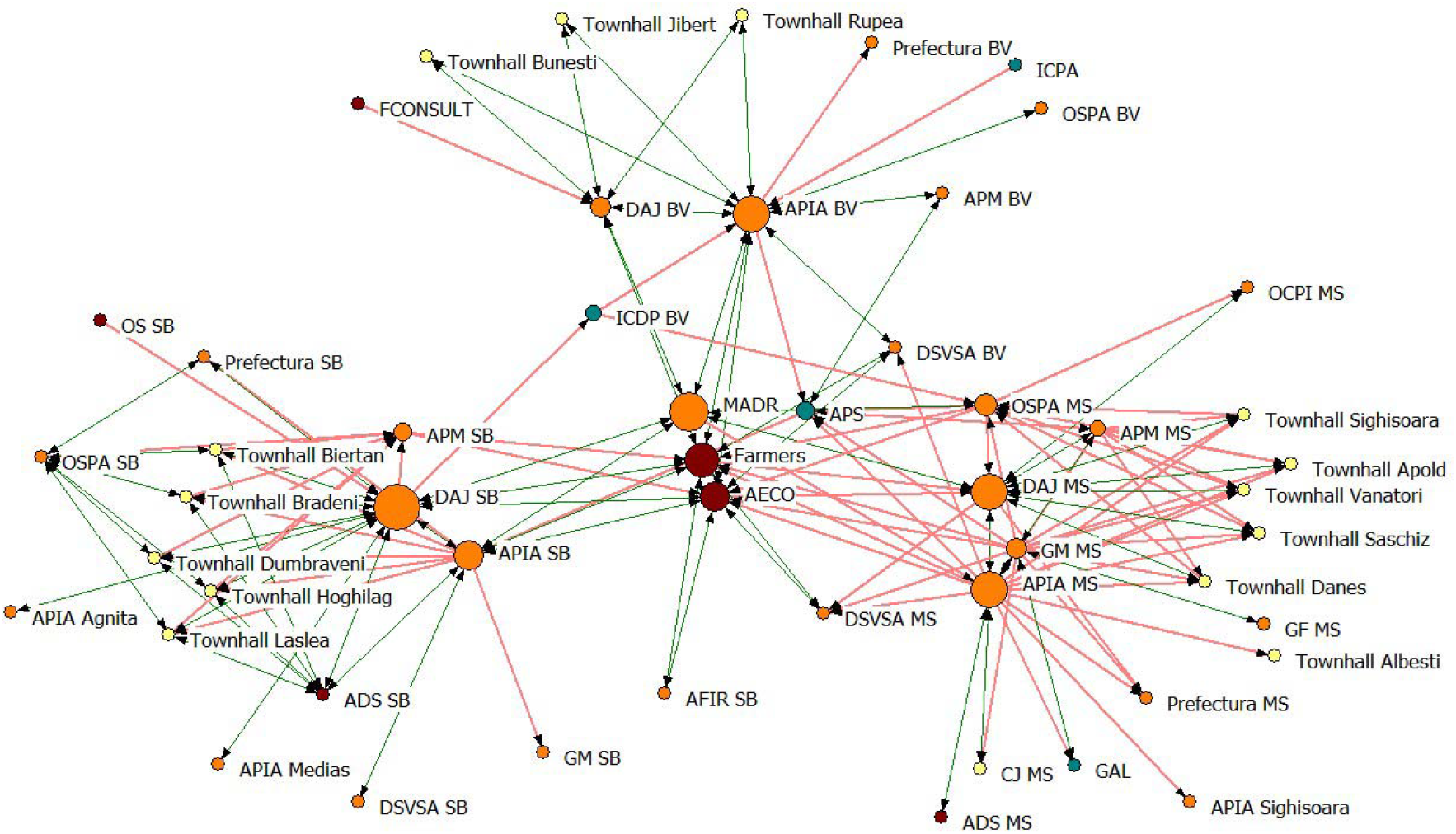
Grassland collaboration network in the Sighisoara area (green lines = reciprocal links of collaboration, red lines = one direction of collaboration, orange circles = public authorities, red circle = enterprises, yellow circle = public administration, green circle = NGO & research & education organizations, size of nodes = betweenness centrality). The full names of organizations are available in Supplementary Data 3.

**Figure 4.**
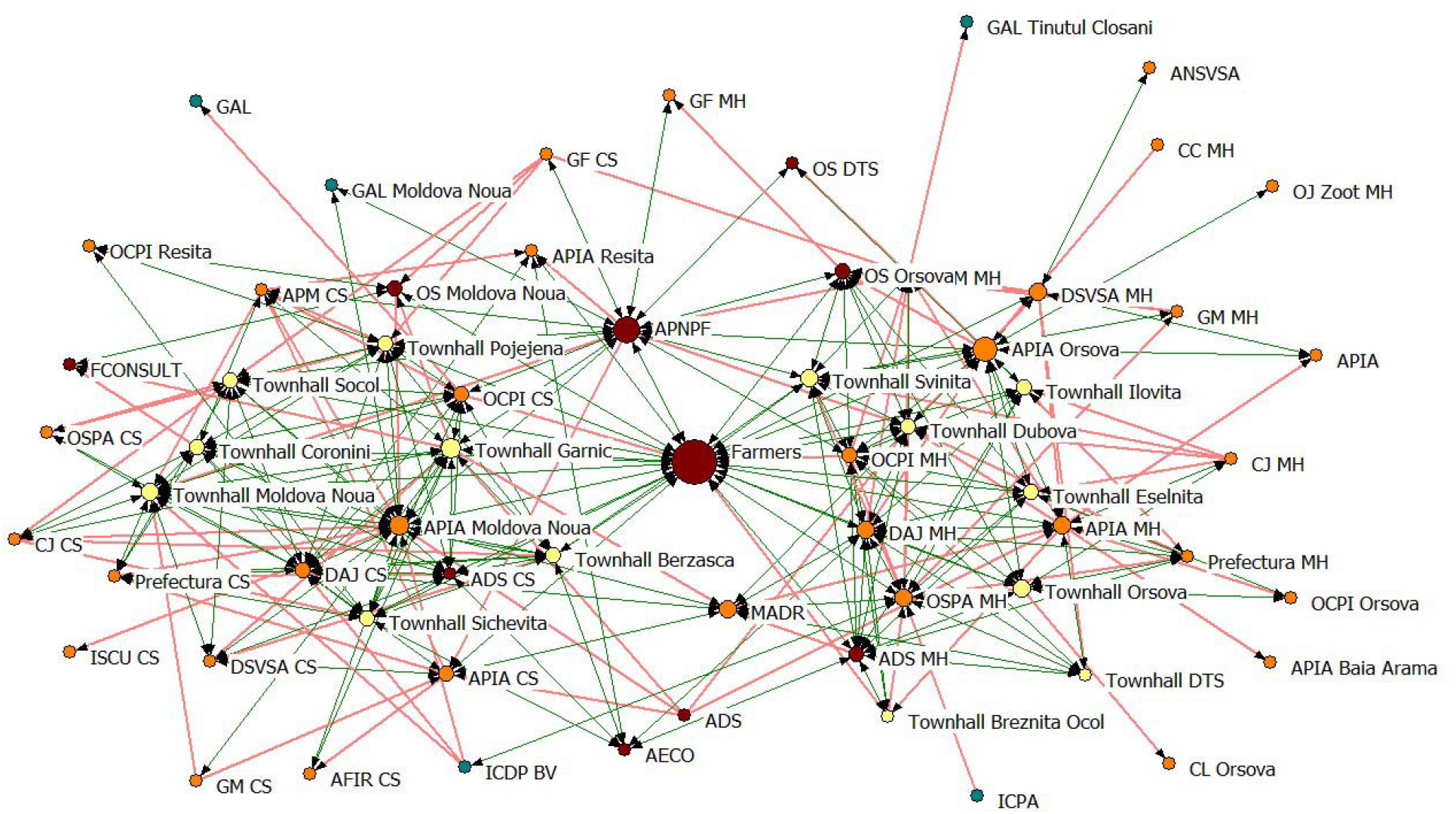
Grassland collaboration network in Iron Gates (green lines = reciprocal links of collaboration, red lines = one direction of collaboration, orange circles = public authorities, red circle = enterprises, yellow circle = public administration, green circle = NGO & research & education organizations, size of nodes = betweenness centrality). The full names of organizations are available in Supplementary Data 3.

When contrasting the in-degree (number of incoming collaborations) and the out-degree centralities (number of outgoing collaborations) with the eigenvector centrality (influential stakeholders in the network) (Figure 5, Supplementary Data 3), the top stakeholders are similar with those resulted from betweenness centrality, with the notable exception of two town halls in Iron Gates network (Moldova Noua and Garnic), involved in advising farmers to comply with the rules for agri-environmental measures.

**Figure 5.**
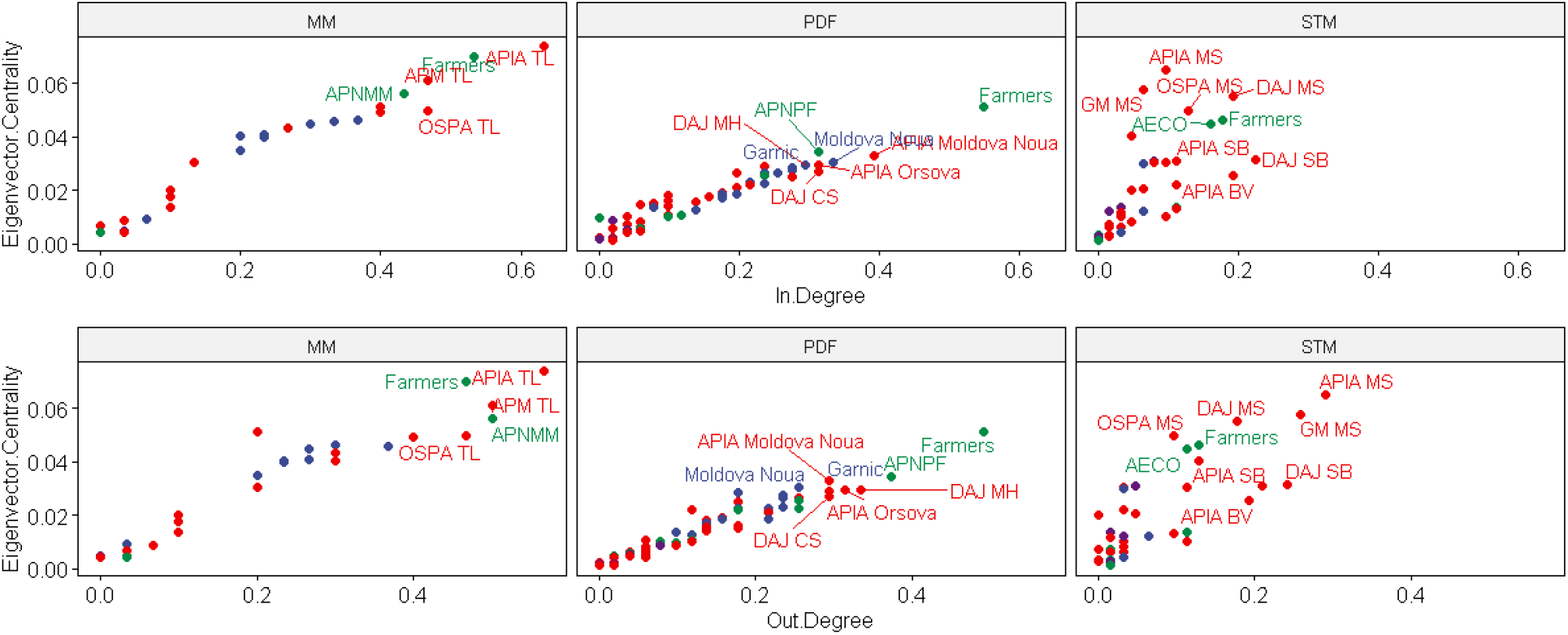
In and out-degree versus eigenvector centrality in grassland collaboration networks in the three protected areas (MM = Macin Mountains; PDF = Iron Gates; STM = Sighisoara). The full names of organizations are available in Supplementary Data 3.

### 3.3. Farmers perception of grassland governance

When analyzing the main issues of grassland governance as perceived by the farmers benefiting of agri-environmental payments resulted that the farmers of the Sighisoara area and Macin Mountains are slightly different from the farmers of Iron Gates on both dimensions of MCA biplot (Figure 6). When interpreting the answers with a contribution to a dimension above the average contributions of all variables in the respective dimension, it resulted that the farmers of the Sighisoara area and Macin Mountains consider that the CAP provides insufficient funds (dimension 1). At the same time, the farmers from Iron Gates consider CAP having a positive influence on grassland management, but there is an issue with late payments. On dimension 2 the pattern indicates that the farmers of Sighisoara and Macin consider CAP as providing a market for grassland products, while the farmers of Iron Gates consider CAP having a positive influence even if the payments are insufficient but too bureaucratic.

**Figure 6.**
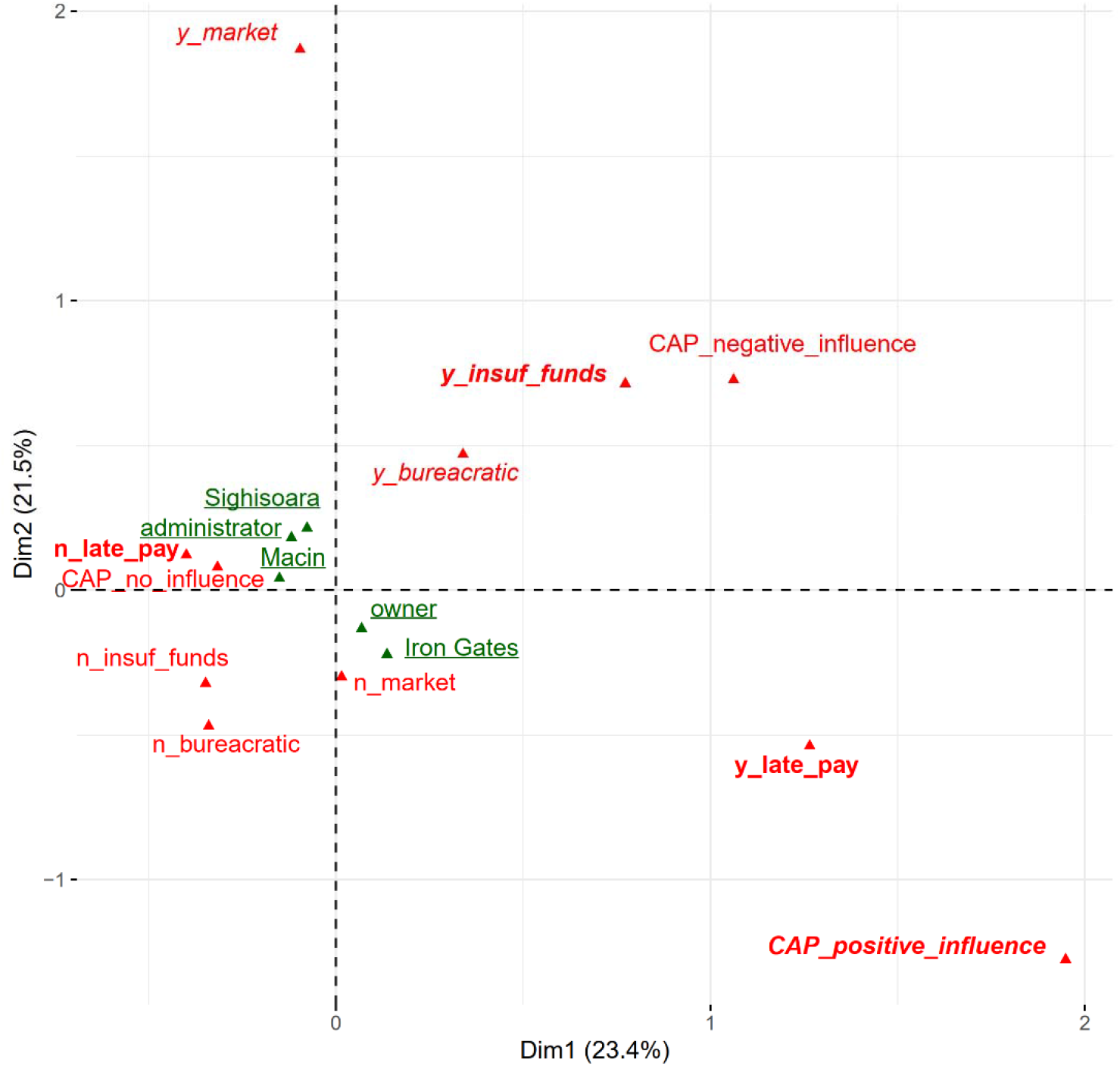
Biplot of MCA on farmers perception of grassland governance (bold = significant on dimension 1, italic = significant on dimension 2, underlined = supplementary variables).

## 4. Discussion

Coordination of networks around grassland management within protected areas with different past management is influenced by the complexity of the administrative institutions at the local level and not by past farming practices and land stewardship. Management actors from areas where the sense of ownership is a long tradition (e.g., Sighisoara area, Figure 3) are not keen to collaborate more than actors from areas with a centralized past, such as Macin Mountains (Figure 2). This can be explained by the fact that Sighisoara area had a large ethnic and social change, with the rapid collapse of the traditional Saxon society and institutions and the formation of new rural communities in many of them being no Saxon ethnics anymore (Fischer et al., 2012; Hartel et al., 2014). In this area, the current rural inhabitants often refer nostalgically to the Saxon society they experienced as being dominated by strong social cohesion, well-defined rules, and communal grassland and forest stewardship forms (Hartel et al., 2014). The political and institutional instability of the recent decades (the communist and post-communist period) does not allow forming of strong social and institutional capitals, the rural communities suffering from rigidity and poverty traps (Mikulcak et al., 2015). Within the above context, the legacy of the past is strong in the people’s collective memory but does not help create a collaborative institutional structure. Institutional networks in all three regions are coordinated by county-level public authorities in charge of agri-environmental payments, which suggest that the EU’s CAP did not genuinely contribute to the formations of farmers’ local associations. The CAP provides additional income for agri-environmental measures undertaken by a group of farmers. However, similar to Germany and Sweden (Leventon et al., 2017), the support is not effective, and the CAP payment criteria should be adjusted to foster better collaboration at local levels.

The number of management actors involved in grassland management is larger in Iron Gates (Figure 4) and Sighisoara area (Figure 3) than in Macin Mountains (Figure 2). The likely reason for this finding is that the region is part of more counties (two in Iron Gates, three in the Sighisoara area compared to one county in Macin Mountains). Local authorities and public administration dominate collaboration networks in all three regions (84% of actors in Macin Mountains, 81% in the Sighisoara area, and 77% in Iron Gates); thus, the size of the network was influenced by administrative jurisdiction and not by diversity of non-public stakeholders (Figures 2–4, Table 2). The importance of inclusion of the local, informal stakeholders and knowledge types into the management of multifunctional landscapes with high socio-cultural and natural values was suggested as a key element for leveraging sustainability transformations responding to future environmental change (Rotherham, 2010; Balázsi et al., 2019). The results of our analyses also reveal a strong relationship between the farmers and the conventional institutional structures involved in the grassland governance and a weak influence of the protected area management institutions on their grassland governance practices (Table 2).

The effect of territorial jurisdiction is also evident when analyzing the mean number of connections of an institution. Institutions participating in grassland management in the Sighisoara area are the least connected (3.98 connection per institution) while in Iron Gates, and Macinului Mountains are well connected (average degree = 7.33 in Iron Gates and 6.58 in Macin Mountains). The actors from the Sighisoara area also display lower reciprocity than the two other networks (65% comparing to over 80%, Table 2). This suggests low levels of bidirectional collaborations between the grassland governance institutions, implying a unidirectional flow of information and power influence within the network. This could represent a challenge in the front of sustainability initiatives, which also implies that the existing grassland governance network is capable to ‘absorb’ genuine information from the outside and turn it into opportunities of grassland management. Indeed, several research initiatives and research communication activities targeting ancient grasslands exist in the Sighisoara region (Hartel et al., 2020); however, virtually no information turns into a real opportunity for the sustainability of these ecosystems.

When interpreting the ERGM results (Table 3 and Supplementary Data 4), the historical management effects on the patterns of relationships in the three protected areas appear to be influenced by the top-down prescriptions of the CAP. The tendency for collaboration between actors is higher than expected in the Sighisoara area and Macin Mountains and without a pattern (higher/lower than expected in Iron Gates (reciprocity effect Table 3). The reciprocity of collaboration in the Sighisoara area is reduced when compared with Iron Gates and Macin Mountains (only 65% compared to 85% in Macin Mountains). However, because the reciprocity is stronger than expected at network-scale effect, there is a high potential for improving if non-public organizations and protected area manager will intensify their trans-counties collaborations (for example, a more active local agro-economy). On the contrary, the higher than expected reciprocity from Macin Mountain suggests that the only way to improve the collaboration is by involving other actors in the network than existing ones (e.g., farmers associations, small enterprises).

In the Sighisoara area, the governance network shows a tendency for sending a request for collaborations to only two other organizations (2-out-star effect positive). However, the effect is reversed at higher orders (e.g., sending a request for collaborations to only three other organizations, 3-out-star negative). The effects for incoming ties (i.e., 2-in-star, 3-out-star) are at random in the Sighisoara area and Macin Mountain, suggesting the lack of popular actors (actors receiving more collaboration requests than expected). Such actors are present in Iron Gates (2-in-star positive), suggesting that high ranking actors in terms of centrality metrics are network leaders.

In the Sighisoara area, the management network also shows a lack of bridging organizations (2-path negative) and a substantial lack of appetence to transfer information to other organizations in the network (AoutS negative). The lack of institutions actively seeking to expand the connections is coupled with avoidance of collaboration between institutions with a similar level of activity (lower or higher) in terms of in-degree and out-degree (AinAoutS negative) complete the picture of an unexpected low collaborative network within Sighisoara area. In the two other networks, most effects in the converged models are at random, with the notable exception of reciprocity (higher than expected) and avoidance of collaboration between institutions with a similar level of activity (AinAoutS negative) and a low degree centralization around popular institutions (2-in-star positive) and a lower chance to break this centralization (A2P-DU negative) in Iron Gates.

The key stakeholders by contribution to information flow in the network (network members with high betweenness centrality, Figure 2–4) are local Agencies for Payments and Intervention for Agriculture in the Sighisoara area and Macin Mountains, and farmers as collective institutions and Iron Gates Natural Park Administration in Iron Gates. From this point of view, CAP contributed to bureaucratic governance in the Sighisoara area (Mikulcak et al. 2013) and Macin Mountains but to a lesser degree in Iron Gates, where the protected area administration is very active. The same pattern is evident when considering eigenvector centrality (higher eigenvector suggesting a higher power of institutions participating in the respective network) and in-degree and out-degree (activity in the network) (Figure 5). In the Sighisoara area and Macin Mountains, the agencies distributing the CAP agri-environmental payments are most active and influential, while in Iron Gates, some farmers, the protected area administrator, and city halls dominate the network.

The history of the investigated protected areas fundamentally influences the farmers and their perceptions and actions on the grassland management (Hanspach et al., 2014); however, our study suggests a diminished influence after only a few years of CAP agri-environmental payments. The MCA results (Figure 6) indicate that the farmers from the Sighisoara area and Macin Mountains perceive the CAP differently than farmers from Iron Gates. Farmers from the Sighisoara area and Macin Mountains have no convergent opinion about CAP influence, while the farmers from Iron Gates (mostly small owners) recognize a positive influence of the CAP. This pattern might be because grasslands from the Sighisoara area and Macin Mountains are lucrative business while in Iron Gates are mostly for subsistence (Manea, 2003; Rey et al., 2007), thus, receiving money from CAP agri-environmental measures is considered by Iron Gates farmers as a good addition to the local economy. The need for additional income s confirmed by the complaining about late payments and insufficient funds. Late payments are not an issue for farmers from the Sighisoara area and Macin Mountains (mostly administrators of leased grassland) but insufficient funds. Notably, in the Sighisoara area and Macin Mountains, farmers consider CAP as providing a market for their products, which is in part because of land concentration in these areas (Calota and Patru-Stupariu, 2019).

Traditionally managed grasslands suffer threats such as intensive, highly specialized management of grasslands and land abandonment (Dorresteijn et al., 2013; Hartel and Plieninger, 2014; Öllerer, 2013). According to the survey results, although farmers receive agri-environmental payments for non-intensive management of their grasslands, they do not consider this as an aid to bring prosperity to rural biodiversity and cultural heritage conservation. Our results showed that in the investigated case studies, CAP and protected area regulations failed to reverse the threats to traditionally managed grasslands and did not foster genuinely new institutional collaborations. Farmers from former state-owned areas are still looking for state advice, while those from areas with a liberal past are not keen to collaborate, and the networks are centralized around regional and local public bodies (Figure 6).

## 5. Conclusions

This research significantly contributes to the consolidation of using social network analysis for understanding landscape governance worldwide (Barnes et al., 2017). Our results provide a solid base for further applications of social science in resource management and to identify key milestones for understanding the evolution of coupled social-ecological systems (Bodin et al., 2019). From the ecological and social point of view, we identified the barriers to the implementation of CAP through the analysis of relevant landscape governance networks in Romania and determined how top-down prescriptions influence network structures. At the European level, the strengthening of the grassland management needs to be deepened, and each strategic plan of the CAP should include a section on how to stimulate knowledge exchange and innovation.

## 6. Acknowledgments

We would like to thank reviewers for their insightful comments on the paper, which helped us to improve the quality of the manuscript. This work was supported by a grant of the Romanian National Authority for Scientific Research (https://uefiscdi.gov.ro), PN-III-P4-ID-PCE-2016-0483.

## Notes

https://github.com/rlaurentiu/grassland_networks

### Competing Interest Statement

The authors have declared no competing interest.

