## Supplementary data 1 for "Governance networks around grasslands with contrasting management history"

General map of Romania, showing the geographic position of the three case studies: Macin Mountains National Park (green); Iron Gates Natural Park (blue); Sighisoara-Tarnava Mare Natura 2000 site (red)

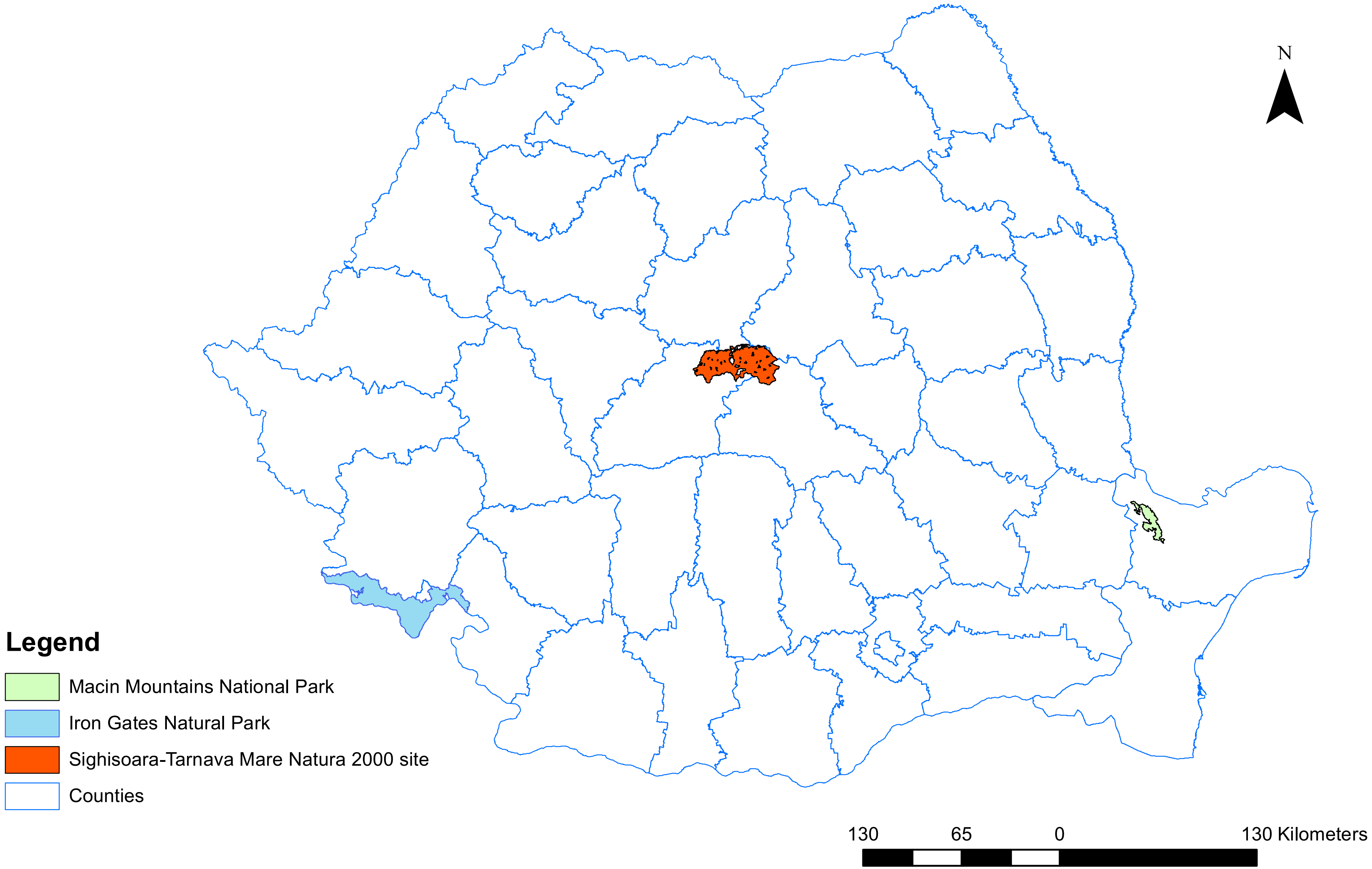

Iron Gates Natural Park

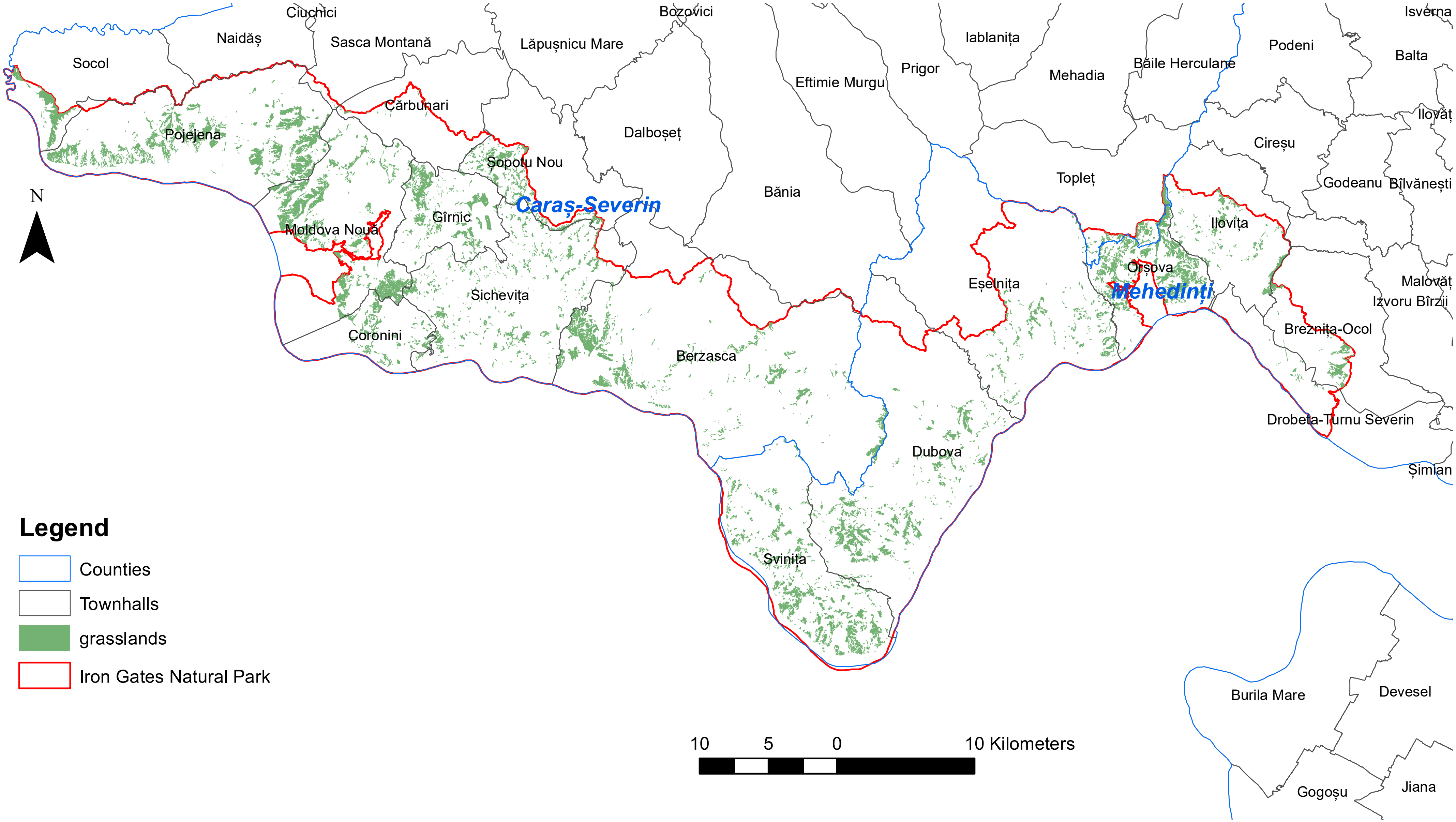

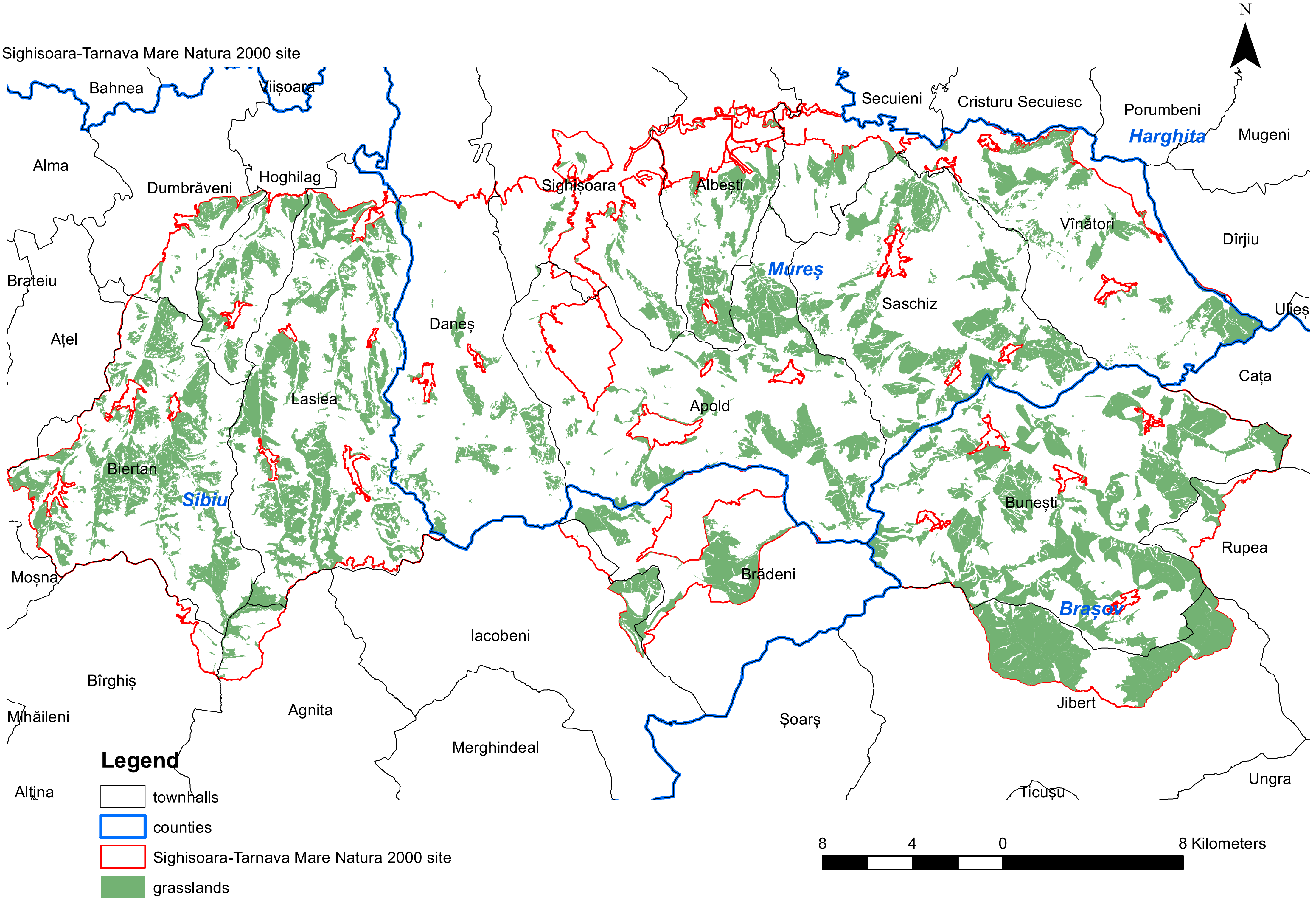

Sighisoara-Tarnava Mare Natura 2000 site
N
Bahnea
Viișoara
Secuieni
Cristuru Secuiesc
Porumbeni
Harghita
Mugeni
Alma
Hoghilag
Dumbrăveni
Sighișoara
Albești
Vînători
Dîrjiu
Brateiu
Mureș
Saschiz
Ulieș
Ațel
Daneș
Cața
Laslea
Apold
Biertan
Sibiu
Bunești
Rupea
Moșna
Brădeni
Brașov
Iacobeni
Jibert
Bîrghiș
Mihăileni
Agnita
Șoarș
Merghindeal
Ungra
Altîna
Ticușu
Legend
townhalls
counties
Sighisoara-Tarnava Mare Natura 2000 site
grasslands
8
4
0
8 Kilometers

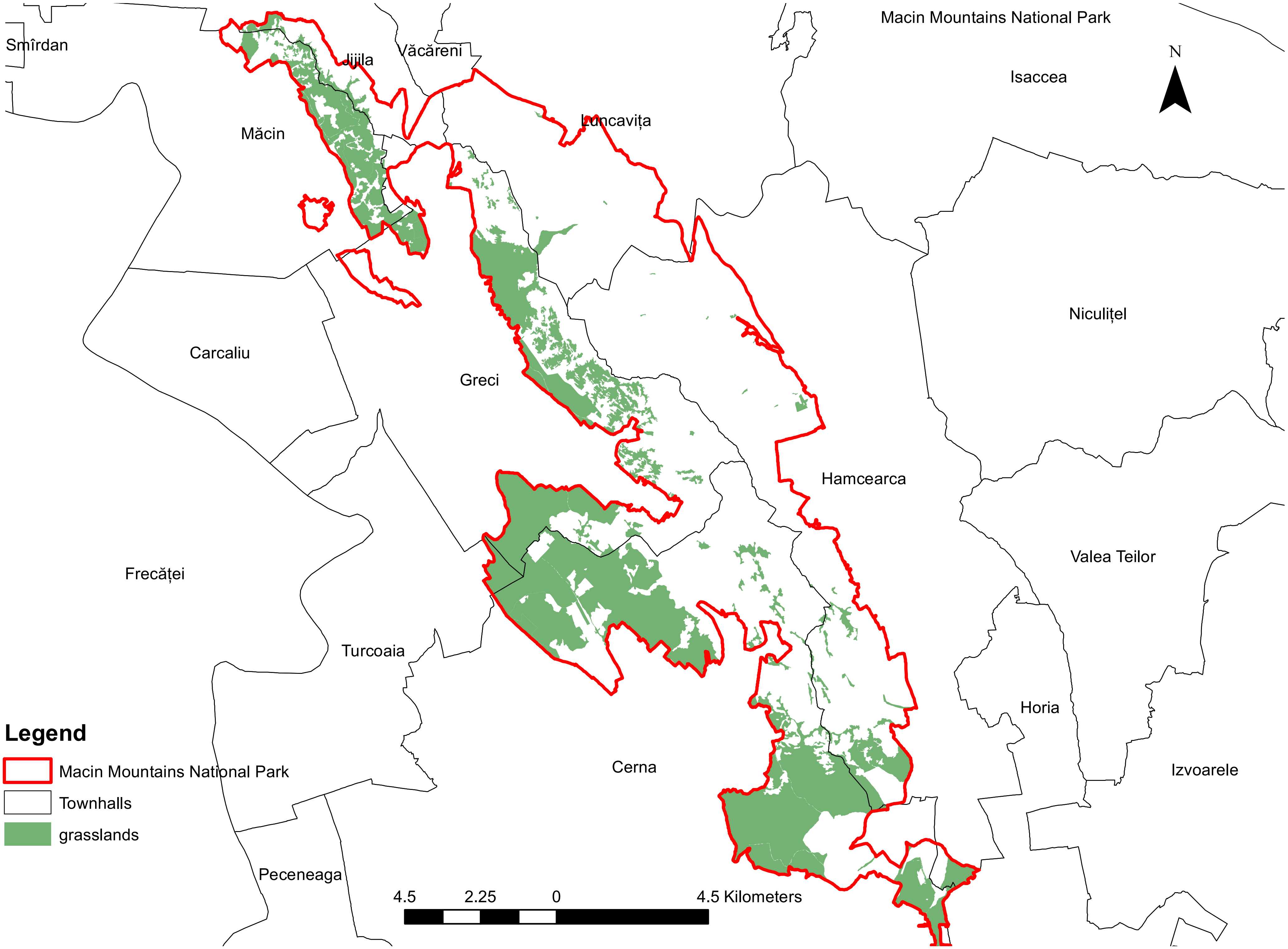

Macin Mountains National Park
N
Smîrdan
Jijila
Văcăreni
Isaccea
Măcin
Luncavița
Niculițel
Carcaliu
Greci
Hamcearca
Frecăței
Valea Teilor
Turcoaia
Horia
Cerna
Izvoarele
Peceneaga
4.5
2.25
0
4.5 Kilometers
Legend
Macin Mountains National Park
Townhalls
grasslands
