## Supplementary data 2 for "Governance networks around grasslands with contrasting management history"

**Supplementary Table 1 - Main characteristics of case study areas**

| Characteristics | Iron Gates | Sighisoara-Tarnava Mare | Macin Mountains |
|---|---|---|---|
| Counties | Caras-Severin, Mehedinti | Sibiu, Mures, Brasov | Tulcea |
| Administrative units | Moldova Nouă, Pojejena, Cărbunari, Coronini, Gârnic, Șopotu Nou, Socol, Sichevița, Berzasca, Orșova, Dubova, Eșelnița, Ilovița, Svinița, Breznița-Ocol | Biertan, Laslea, Brădeni, Dumbrăveni, Agnita, Hoghilag, Daneș, Apold, Albești,Vânători, Saschiz, Sighișoara, Bunești, Jibert | Măcin, Isaccea, I.C. Brătianu, Jijila, Văcăreni, Luncavița, Niculițel, Frecăței, Nalbant, Izvoarele, Valea Teilor, Horia, Hamcearca, Cerna, Turcoaia, Greci, Carcaliu, Smârdan |
| Main protected areas | Iron Gates Natural Park (category V IUCN and Natura 2000 site) | ROSCI0227 Sighisoara-Tarnava Mare | Macin Mountains National Park (category II IUCN and Natura 2000 site) |
| Geographical units | Locvei, Almajului and Mehedinti Mountains, Mehedinti Tableland | Tarnavelor and Hartibaciului Tablelands | Macin Mountains |
| Biogeographical region | Continental | Continental | Steppic |
| Altitude (min-max-med, meters) | 28-972-368 | 315-829-541 | 4-466-214 |
| Climate | Hot-summer humid continental | Warm-summer humid continental | Hot-summer humid continental & warm-summer humid continental |
| Main grassland types | Subcontinental peri-Pannonic scrub, Rupicolous calcareous or basophilic grasslands of the Alysso-Sedion albi, Xeric sand calcareous grasslands, Rupicolous pannonic grasslands (Stipo-Festucetalia pallentis), Semi-natural dry grasslands and scrubland facies on calcareous substrates (Festuco-Brometalia), Hydrophilous tall herb fringe communities of plains and of the montane to alpine levels | Semi-natural dry grasslands and scrubland facies on calcareous substrates (Festuco-Brometalia), Sub-Pannonic steppic grasslands, Lowland hay meadows (Alopecurus pratensis, Sanguisorba officinalis), Hydrophilous tall herb fringe communities of plains and of the montane to alpine levels | Ponto-Sarmatic steppes, Ponto-Sarmatic deciduous thickets, Pannonic salt steppes and salt marshes |
| Grassland surface | 18% of Iron Gates Natural Park | 26% of ROSCI0227 Sighisoara-Tarnava Mare | 18% of Macin Mountains National Park |
| Past ownership type | Small private farms | mixture of communal, collective, and private small farms ( | Large collective farms, state-run management regime |
| Present ownership type | Small private farms | Large and small private farms | Large and small private farms |
